# Carnosinylation of Cardiac Antigens Attenuates Immunogenic Responses and Improves Function in Failing Hearts

**DOI:** 10.1101/2025.08.22.671840

**Authors:** Benjamin Doelling, Mamata Chaudhari, David Hoetker, Kenneth Brittian, Yibing Nong, Jonah K Stephan, Ibrahim Jouja, Thomas Mitchell, Marcin Wysoczynski, Aruni Bhatnagar, Steven P Jones, Shahid P Baba

## Abstract

**Objective:** To investigate the effects of carnosine on heart failure and to examine whether this is associated with reduced immunogenicity of oxidatively-generated aldehyde modified proteins.

**Background:** Heart failure is associated with the accumulation of lipid derived aldehydes that form immunogenic protein adducts. However, the pathological impact of these aldehydes and aldehyde-modified proteins in heart failure has not been assessed. Histidyl dipeptides, such as carnosine found in the heart, bind to aldehydes, and their protein adducts. However, the effects of carnosine on heart failure or the antigenicity of aldehyde modified proteins have not been studied.

**Methods:** Male, wild type C57BL/6J mice were subjected to either sham or transverse aortic constriction (TAC) surgery. To increase carnosine levels, they were placed on drinking water with or without β-alanine prior to surgery, and for the remainder of the study. Cardiac function was evaluated by echocardiography, and the levels of histidyl dipeptides, immune cell populations, and CD4^+^ T cell activation were assessed via LC-MS/MS and flow cytometry, respectively.

**Results:** Myocardial levels of histidyl dipeptides decreased at both 3- and 8-weeks post-TAC. Supplementation with β-alanine increased myocardial histidyl dipeptide levels, attenuated adverse cardiac remodeling, and reduced aldehyde stress. Carnosine formed covalent bond with protein-bound aldehydes in the failing heart, reducing their antigenic potential and decreasing activation of dendritic cells and CD4^+^ T cells *in vitro*. β-alanine supplementation decreased the population of CD11b^+^CD64^-^Ly6G^+^ neutrophils and CD4^+^ CD44^+^ effector T cells in the failing heart.

**Conclusions:** Increasing myocardial carnosine levels reduces aldehyde stress, dampens maladaptive immune responses, and preserves cardiac function during heart failure.

**HIGHLIGHTS:**

- Levels of endogenous dipeptide carnosine are depleted in failing hearts, while supplementation of the carnosine precurson β-alanine increases myocardial carnosine and preserves cardiac function during heart failure.
- Heart failure is associated with increased activation and infiltration of CD4^+^ T cells and generation of aldehyde modified protein adducts in failing hearts.
- The free aldehyde moiety of aldehyde modified protein adducts activates CD4^+^ T cells through dendritic cell presentation and capping this moiety with carnosine diminishes their antigencity.
- Increasing myocardial carnosine levels diminishes aldehyde stress and activation of CD4^+^ T cells during heart failure.

**GRAPHICAL ABSTRACT:** 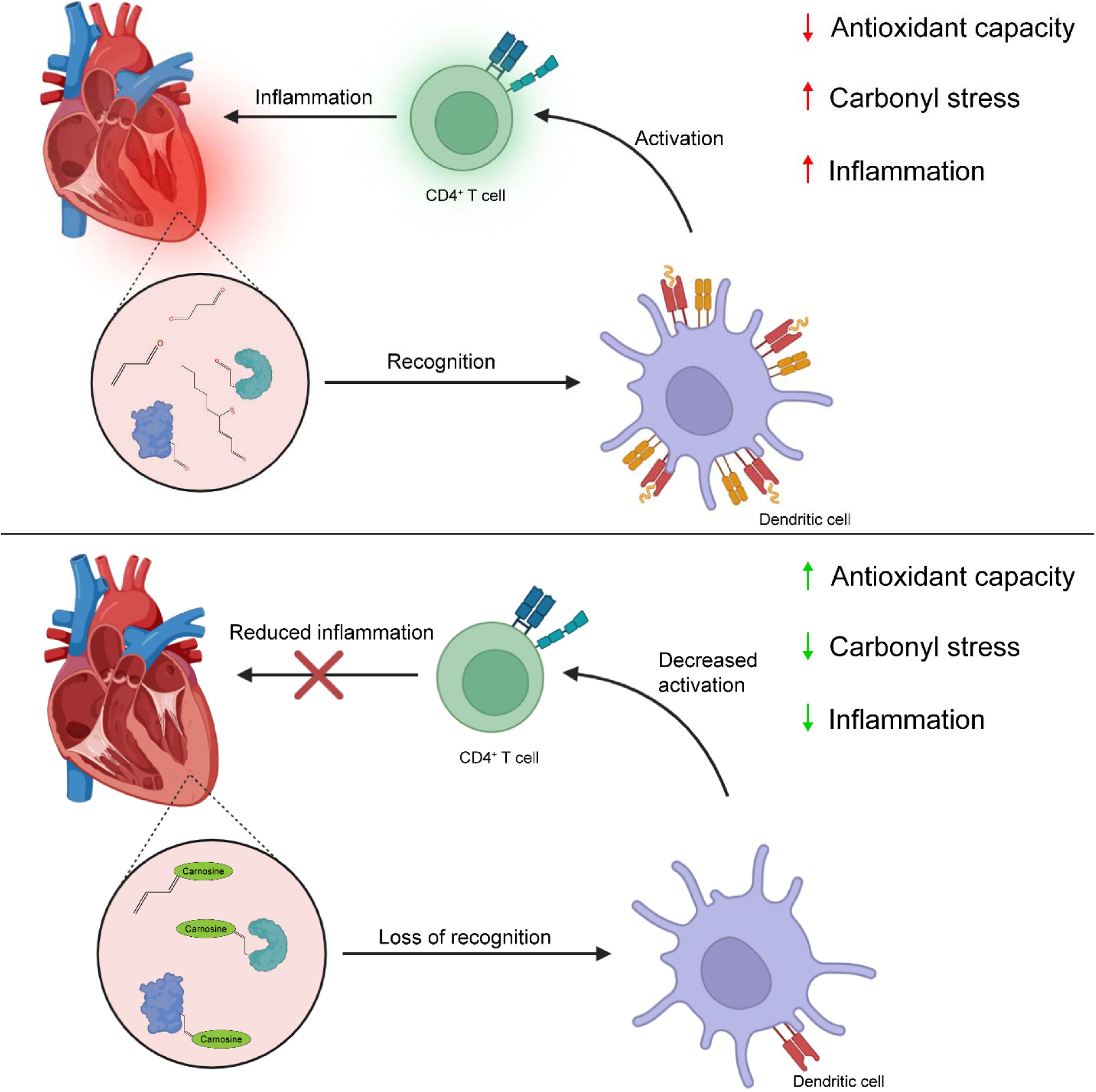

## INTRODUCTION

Heart failure is a complex pathological state which involves progressive cardiac dysfunction and is associated with significant increases in both - oxidative stress and inflammation^1–4^. Although oxidative injury and tissue-damaging inflammation have been shown to be a prominent features of the failing heart, indiscriminate immunosuppression and antioxidant therapies have not been shown to be effective, and in some cases, may even worsen heart failure by blocking necessary reparative processes^5–7^. Effective and feasible strategies to target inflammation and oxidative stress are urgently needed but require a deeper understanding of the cellular and molecular processes that contribute to the pathophysiology of heart failure.

The immune response observed in the failing heart is, in part, triggered by the release of autoantigens which instigate non-resolving adaptive immune responses^8,9^. They can be generated by oxidative stress and by binding highly toxic lipid peroxidation derived carbonyls, such as acrolein and 4-hydroxynonenal, to proteins^10,11^. These carbonyls are the downstream effectors of oxidative stress that possess two highly electrophilic groups - an α,β-unsaturated motif and a carbonyl group. Because of this structural motif, they react readily with nucleophilic side chains of proteins, particularly histidine and lysine^12,13^. Importantly, the remaining free electrophilic group on these aldehyde-modified proteins causes them to function as aldehyde modified damage associated molecular patterns (ADAMPs) that instigate both innate and adaptive immune responses^14^. Although several studies have shown that ADAMPs are generated in the heart ^15,16^, and it has been speculated that they contribute to the pathogenesis of heart failure, there is no direct evidence linking ADAMPs to inflammation in the context of heart failure. Given that reactive aldehydes accumulate in the failing heart, we posit that ADAMPs instigate inflammation in a pressure overload model of heart failure.

Reactive aldehydes in the heart are metabolized through several enzymatic pathways, such as aldehyde dehydrogenase^17^, aldo-keto reductases^18^, and by conjugation with reduced glutathione^19,20^, catalyzed by glutathione-S-transferases. In addition to these pathways, reactive aldehydes are removed by forming conjugates with histidyl dipeptides, such as carnosine (β-alanyl-L-histidine) and anserine (β-alanyl-N^π^-methylhistidine), which are synthesized via the enzyme carnosine synthase (CARNS) and carnosine methyl transferase respectively^21,22^. Endogenously carnosine levels in both skeletal muscle^23^ and heart^24^, can be increased by supplementation with β-alanine, which leads to the clearance of reactive aldehydes from these tissues^25,26^. Previous studies have shown that an increase in carnosine levels can diminish aldehyde stress during strenuous exercise^27^, protect heart from ischemia reperfusion injury^24^, attenuate atherosclerosis^28^, and improve the glucose control in prediabetic and type 2 diabetes patients^29^. Nevertheless, upon binding with the reactive aldehydes, carnosine-aldehyde conjugates still retain a free electrophilic moiety that enables them to form a covalent bond with the nucleophilic amino acids of proteins, resulting in the generation of carnosinylated proteins^30^. While we have identified such carnosinylated proteins in aging hearts^30^, it remains unknown whether carnosine could bind with the free aldehydic moiety of ADAMPs, result in their carnosinylation, neutralize their immunogenicity, and influence the inflammatory response and cardiac function during heart failure.

In the present study, we tested the hypothesis that increasing the levels of myocardial carnosine in failing hearts will sequester reactive aldehydes, carnosinylate and diminish the antigenicity of ADAMPs, and consequently attenuate cardiac inflammation and function. We show that myocardial histidyl dipeptides are decreased in failing hearts, β-alanine supplementation increases carnosine levels and sequesters reactive aldehydes, preserves cardiac function, and reduces ADAMPs antigenicity. Further, we show that “capping” of ADAMPs by carnosine prevents CD4^+^ T cell activation, thus suggesting that this posttranslational modification may attenuate immune-mediated injury in the failing heart.

## MATERIALS AND METHODS

### Animals

Wild-type C57BL/6J and B6N.B6-Tg (Nr4a1-EGFP/cre) 820Khog/J (Nur77-GFP) mice were obtained from The Jackson Laboratory (Bar Harbor, ME). Mice were treated in accordance with the Guide for the Care and Use of Laboratory Animals (Institute of Laboratory Animal Resources, 2011) as adopted and promulgated by the National Institutes of Health. All animal experiments were performed under protocols approved by the Institutional Animal Care and Use Committee of the University of Louisville.

### Transverse aortic constriction surgery and β-alanine treatment

C57BL/6J mice, 12–15 weeks old, male, were either pretreated with drinking water alone or drinking water supplemented with β-alanine (20 g/L) for 7 days and then subjected to sham and transverse aortic constriction (TAC) surgeries as described^18,31–33^. Supplementation was continued for the duration of the experiment, with water changing weekly. Briefly, following anesthesia (i.p. 50 mg/kg sodium pentobarbital and 50 mg/kg ketamine hydrochloride), the mice were orally intubated with polyethylene-60 tubing; and ventilated with a mouse ventilator (Hugo-SACS ELEKTRONIK). Analgesia (meloxicam, 20 mg/kg) was provided subcutaneously prior to surgery and 24 and 48 h post-surgery, as relevant. Prior to skin incisions, mice were also given local anesthetic (bupivacaine liposomal suspension, 5.3 mg/kg). The aorta was visualized following an intercostal incision. A 7–0 nylon suture was used to loop around the aorta between the brachiocephalic and left common carotid arteries. The suture was tied around a blunted 27-gauge needle placed adjacent to the aorta to constrict the aorta to a reproducible diameter. The needle was removed, and the chest was closed in layers. The mice were extubated upon recovery from spontaneous breathing. The surgeon was blinded to mouse genotype and animal group assignment. Sham mice were subjected to the same procedure as the TAC group, except that the suture was not tied. Mice were followed for the indicated durations.

### Echocardiography

Transthoracic echocardiography of the left ventricle was performed using a VisualSonics Vevo 3100 system equipped with a MX550D probe as described^31,33–35^. Serial echocardiograms were obtained at 4- and 8-weeks post-TAC. Briefly, the mice were anesthetized with 2% isoflurane, body temperature was maintained at 37.0 ± 0.5°C throughout the procedure. We obtained 2D images of parasternal long-axis, short-axis, apical 4-chamber and aortic arch views. End systolic and diastolic volumes were determined from long-axis B-mode views, and ejection fraction was calculated as [(stroke volume)/(end diastolic volume)]*100%. Left ventricular inner diameter during diastole (LVIDd), left ventricular inner diameter during systole (LVIDs), and heart rate were determined from short-axis M-mode images. The sonographer was blinded to animal group assignments. The peak blood velocities before the point of constriction (V0) and at the point of constriction (V1) were measured via pulse-wave Doppler at the aortic arch view 8 weeks post-TAC. The estimated trans-TAC pressure gradient (ΔP) was then calculated using the Bernoulli equation ΔP=4*(V1^2^-V0^2^), where velocities are in m/s and ΔP in mmHg. TAC mice with pressure gradient less than 40 mmHg were excluded, leading to exclusion of 2 mice from the β-alanine supplemented TAC mice in Figure 2. There were no significant differences in ΔP between the supplemented and non-supplemented mice after exclusion.

### Measurement of histidyl dipeptides by mass spectrometry

Levels of histidyl dipeptides in the heart were determined using UPLC-ESI-MS/MS as described^24,25^. Briefly, pulverized cardiac tissue was homogenized in a solution of 10 mM HCl containing 5 μM carnosine-d_4_ and 5 μM anserine-d_4_ as internal standards. Homogenates were sonicated on ice, centrifuged, and collected supernatants were diluted 100x in a solution of 75:25 acetonitrile:water. 5 μL of sample was injected into a Waters ACQUITY UPLC H-Class system equipped with a Waters Acquity BEH HILIC column. Analytes were eluted using a binary solvent system consisting of 95% water, 5% acetonitrile, 10 mM ammonium formate, 0.125% formic acid (HILIC A) and 5% water, 95% acetonitrile, 10 mM ammonium formate, 0.125% formic acid (HILIC B). Elution began with 0.1:99.9 A:B at 0.55 mL/min, changing to 99.9:0.1 A:B over the course of 5 min followed by return to 0.1:99.9 A:B over 0.5 min. Eluates were then analyzed using a Xevo TQ-S micro triple quadrupole mass spectrometer. Transition ions used for quantification were as follows: Carnosine, m/z 227 → 110; carnosine-propanal, m/z 283 → 166; carnosine-propanol, m/z 285 →110; carnosine-d_4_, m/z 231 → 110; N-acetylcarnosine, m/z 296 → 110; anserine, m/z 241 → 109; anserine-d_4_ 245 → 109. Dipeptides were quantified by comparing peak areas to peak areas of the appropriate internal standard and normalized to protein concentration.

### Quantification of protein carbonyl content

Cardiac lysates were evaluated for carbonyl content using a 2,4-dinitrophenylhydrazine (DNPH) based colorimetric assay^36,37^. Heart tissues were homogenized in RIPA buffer, sonicated on ice, and centrifuged. Protein concentration was determined by Lowry assay and samples were diluted to 1 mg/mL in diH_2_O. 600 μL of diluted sample was incubated with 120 μL of 10 mM DNPH for 1 h at room temperature (RT), followed by an addition of 720 μL ice-cold 20% trichloroacetic acid (TCA) and 15 min incubation on ice. Samples were centrifuged, supernatant was discarded and pellet washed with 20% TCA and centrifuged again. Supernatant was then discarded, and samples were repeatedly washed with 1 mL 1:1 ethyl acetate:ethanol and centrifuged for a total of 3 washes. After the final centrifugation, pellets were dried and resuspended in 600 μL of 6 M guanidine-HCl. Absorbance at 400 nm was read, using 6 M guanidine-HCl as a blank. The protein carbonyl content was determined using the following formula:

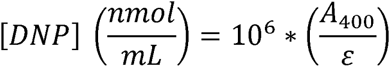

where A_400_ is blank-corrected absorbance at 400 nm and ε = 22,000 M^-1^cm^-1^, the molar extinction coefficient for DNP. Carbonyl concentration was normalized to protein concentration.

### Generation of aldehydes and aldehyde-modified proteins

Malondialdehyde was prepared by incubating 500 mM 1,1,3,3-tetramethoxypropane in 1 M HCl at 37°C for 10 min followed by a 1:19 dilution in 100 mM sodium phosphate buffer (pH 7.4) to quench the reaction^38^. Acrolein was prepared by incubating 250 mM acrolein dimethyl acetal in 100 mM HCl at 37°C for 1 h followed by a 1:9 dilution in 100 mM sodium phosphate buffer (pH 7.4) to quench the reaction^39^. 4-hydroxy-2-nonenal (HNE) was purchased from Abcam and diluted to appropriate stock concentrations in 100 mM sodium phosphate buffer (pH 7.4). Hen egg lysozyme (HEL) was purchased from Roche and prepared at 2 mg/mL in 100 mM sodium phosphate buffer (pH 7.4). Cardiac tissue was homogenized in 100 mM sodium phosphate buffer (pH 7.4), followed by sonication and centrifugation. Cardiac protein lysate diluted to 2 mg/mL in 100 mM sodium phosphate buffer (pH 7.4) and hen lysozyme (HEL) were incubated with 1 mM of either MDA, acrolein, or HNE at 37°C for 1 h. Following incubation, the samples were filtered using Amicon Ultra 3 kDa MWCO centrifugal filters (Sigma-Aldrich) to remove unreacted aldehyde. The protein-aldehyde adducts (2 mg/mL) were further incubated with either 10 mM carnosine at 37°C or reduced with 10 mM NaBH_4_ at RT for 1 h and filtered using 3 kDa MWCO centrifugal filters.

### Western blotting analysis

Cardiac lysates from sham and TAC hearts were homogenized 10% (w/v) in RIPA buffer: 50 mM Tris-HCl, 150 mM NaCl, 1 mM EDTA, 0.25% sodium deoxycholate, 1% Triton X-100, 0.1% SDS with 1% Halt protease/phosphatase inhibitor (Thermo Scientific). Lysates were centrifuged and the supernatants were analyzed via Western blotting as described previously^40^. Similarly, HEL modified with acrolein or HNE, both with and without carnosine, were immunoblotted and developed using anti-acrolein (1:1000), anti-HNE (1:1000), and anti-carnosine (1:1000) antibodies as noted. The immunoblot images were captured using a Bio-Rad ChemiDoc MP imaging system. Band intensity was quantified using ImageJ software and normalized to amido black total protein staining.

### Generation and culture of bone marrow-derived dendritic cells

Bone marrow-derived dendritic cells (BMDCs) were prepared by incubation of bone marrow myeloid precursor cells with the growth factor granulocyte-monocyte colony-stimulating factor (GM-CSF) as described before^41,42^. Briefly bone marrow was isolated by flushing the tibias and femurs of male, wild type C57BL/6J mice with RPMI-1640. Single cell suspensions were generated by passing the bone marrow suspension through a 25 G needle, followed by a 70 μm filter. Cells were plated on untreated 10 cm plates at 2×10^5^ nucleated cells/mL in BMDC media: RPMI-1640 containing 10% heat inactivated FBS, 1% penicillin/streptomycin, 50 μM β-mercaptoethanol, and 40 ng/mL GM-CSF. Additional BMDC media was replaced on day 3, and half of the media was replaced with fresh media on day 6 post-plating. On day 8, non-adherent and loosely adherent cells were collected and plated for further experiments. These populations were verified as 80%+ CD11c^+^ by flow cytometry and were considered bone marrow derived dendritic cells (BMDCs). For activation experiments, BMDCs were plated on nontreated 12-well plates at 2×10^5^ cells/mL in BMDC media and incubated with 100 mg/well of either HEL, MDA-modified HEL, carnosinylated HEL-MDA, or NaBH_4_ reduced HEL-MDA for 6 hours. Cells were collected and stained for surface expression of CD45, CD11b, CD11c, CD80, CD86, and MHCII (I-A/I-E). Flow cytometry data was acquired on a BD LSRFortessa X-20 cytometer and data was analyzed using FlowJo software.

### Splenocyte culture and activation

Spleens from both WT C57BL/6J and Nur77-GFP mice were removed and placed in Hanks’s buffered salt solution (HBSS) on ice. Splenocytes were liberated by grinding the spleen against a 70 μm strainer using the rubber end of a 5 mL syringe plunger. The cells were pelleted by centrifugation at 500xg for 5 min, then the pellet was resuspended in 5 mL of RBC lysis buffer (BioLegend) for 3 min at RT. Lysis was halted by adding 15 mL HBSS and the suspension was centrifuged as before. The cell pellet was resuspended and nucleated cells were counted and plated on non-treated 12-well plates at 5×10^5^ cells/mL in splenocyte media: RPMI-1640 containing 10% heat inactivated FBS, 1% penicillin/streptomycin, and 50 μM β-mercaptoethanol. For HEL experiments, cells were treated with 200 μg/well of HEL, HEL-MDA, carnosinylated HEL-MDA, or NaBH_4_ reduced HEL-MDA and allowed to incubate for 72 h. For cardiac lysate experiments, cells were treated with 200 μg/well of lysate, aldehyde-modified lysate, carnosinylated aldehyde-lysate, or NaBH_4_ reduced aldehyde-lysate for 72 h. Cells were then collected and stained for surface expression of CD45, CD3e, CD4, and CD44 (see Supplemental Table I for details). Flow cytometry data was acquired on a BD LSRFortessa X-20 cytometer and data was analyzed using FlowJo software.

### Quantitative flow cytometry of cardiac tissues and mediastinal lymph nodes

WT C57/BL/6J mice were subjected to sham and TAC surgeries and pretreated with water alone or β-alanine in water as described above. Hearts from the sham and TAC operated mice were isolated, cannulated through the aorta, and the coronary circulation was perfused with 37°C HBSS for 5 min. The atria were removed, and the ventricles were finely minced using a sterile razor blade. Minced hearts were incubated in HBSS containing 1000 U/mL Collagenase II (Worthington Biochemical) for 45 min at 37°C with occasional trituration using a serological pipette^43^. Digested hearts were passed through a 40 μm filter to create a single cell suspension. Isolated cells were centrifuged and resuspended in Cell Staining Buffer (CSB, BioLegend) and nucleated cells were counted using an acetic acid/methylene blue solution (Stemcell Technologies). Cells were liberated from isolated mediastinal lymph nodes (mLNs) by grinding the spleen in CSB against a 70 μm strainer using the rubber end of a 5 mL syringe plunger. Cells from mLNs were counted using a Countess II automated cell counter (Thermo Scientific). The cells were stained by incubation with an antibody cocktail diluted in CSB for 20 min in the dark at RT followed by two washes with CSB. Cells were stained for surface expression of CD45, CD11b, Ly6G, CD64, CD3e, CD4, CD8a, and CD44. Flow cytometry data was acquired on a BD LSRFortessa X-20 cytometer and data was analyzed using FlowJo software v.10.10.0 (BD Biosciences).

### Real-time quantitative PCR

Total RNA was isolated from heart tissues using the RNeasy Fibrous Tissue Mini Kit (Qiagen). RNA concentration and quality were determined using a NanoDrop 2000c Spectrophotometer (Thermo Scientific). Real-time quantitative PCR (RT-qPCR) was performed as described previously^18,31^. Briefly, cDNA was synthesized using the SuperScript IV VILO kit (Invitrogen) with 1000 ng RNA as input. PCR reactions were carried out in a 384-well plate using PowerUp SYBR Green Master Mix (Applied Biosystems) and RT-qPCR was performed on a QuantStudio 5 System (Applied Biosystems). Relative gene expression was normalized to the housekeeping gene *Hprt* and calculated using the 2^-ΔΔCT^ method.

### Statistical analysis

Data are presented as mean ± SEM unless otherwise noted. For single variable comparisons between two groups, p value was determined using Student’s t test. For single variable comparisons between more than two groups, p value was determined using one-way analysis of variance (ANOVA) with Sidak’s *post hoc* correction for multiple comparisons. For echocardiogram data, p values were determined using repeated measures ANOVA with Fisher’s LSD test for comparisons between supplemented and non-supplemented groups at each time point. Outliers within a sample group were detected using Grubbs’ Test with an alpha value of 0.05, resulting in removal of a single data point in Figure 1 and a single data point in Figure 7. Statistical analysis was performed using Prism software v. 10.3.0 (GraphPad Software).

**Figure 1.**
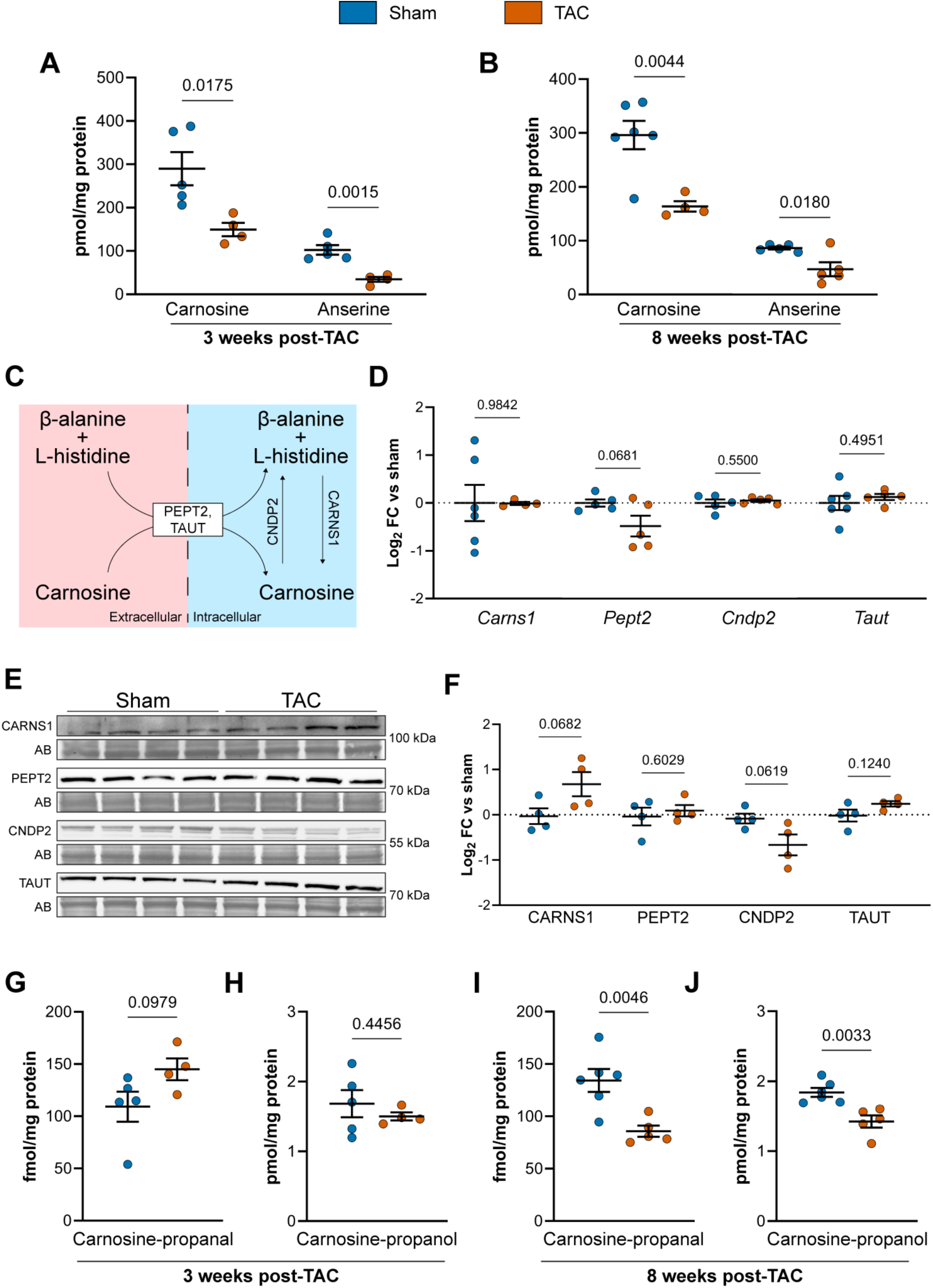
Histidyl dipeptides are decreased in failing hearts. Male wild type C57BL/6J 12-week-old mice were subjected to sham or transverse aortic constriction (TAC) surgery. Myocardial levels of carnosine and anserine were measured by LC-MS/MS using carnosine-d4 as internal standard (IS) after **(A)** 3 and **(B)** 8 weeks of sham and TAC surgery. **(C)** Schematic showing enzymes and transporters involved in carnosine homeostasis. Changes in the expression of carnosine synthase (CARNS1), carnosinase 2 (CNDP2), peptide transporter 2 (PEPT2) and taurine transporter (TAUT) were measured 3 weeks post-surgery at the **(D)** transcript and **(E, F)** protein level using RT-qPCR and Western blotting respectively. **(G)** Levels of carnosine-propanal and carnosine propanol were measured in the sham and TAC operated hearts by LC/MS/MS using carnosine-d_4_ as an internal standard after **(G)** 3 and **(H)** 8 weeks of sham and TAC surgery. Data shown as mean ± SEM, p-values determined using Student’s t-test, n = 4 - 5 mice per group.

**Figure 2.**
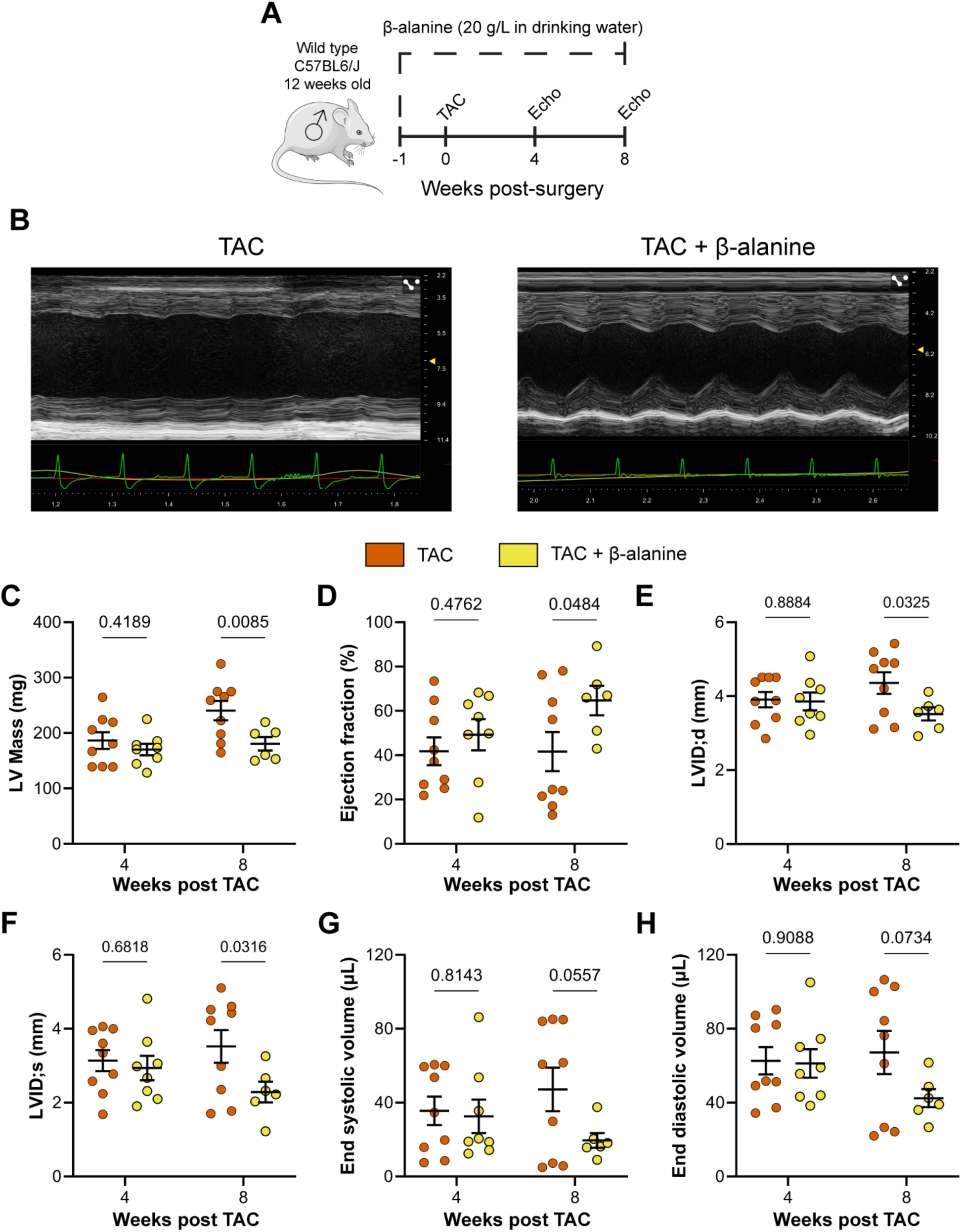
β-alanine supplementation prior to transverse aortic constriction (TAC) attenuates cardiac remodeling. **(A)** Male wild type male C57BL/6J were supplemented with either drinking water alone or drinking water with β-alanine (20 g/L) one week before TAC surgery. β-alanine supplementation was continued until the end of the experiment. After 4- and 8-weeks post-surgery cardiac parameters were evaluated via echocardiography. **(B)** Representative M-modes images, **(C)** left ventricular (LV) mass, **(D)** ejection fraction, **(E)** LV internal diameter diastole (LVID;d), **(F)** LV internal diameter systole (LVID; s), **(G)** end systolic volume, and **(H)** end diastolic volume. Data shown as mean ± SEM, p-values determined using Student’s t-test or repeated measures ANOVA as appropriate, n = 6 - 9 mice per group.

## RESULTS

### Myocardial histidyl dipeptide homeostasis is disturbed in the failing heart

We previously showed that myocardial histidyl peptides, including carnosine (β-alanyl-L-histidine) and anserine (β-alanyl-N^π^-methyl-L-histidine), are decreased in failing hearts^44^; however, the specific stage of heart failure at which this depletion occurs, as well as the underlying mechanisms remained unclear. To determine when histidyl dipeptides are reduced during the progression of heart failure, we subjected wild-type (WT) male C57BL/6J mice to either sham or transverse aortic constriction (TAC) surgery and measured myocardial histidyl dipeptide levels at 3- and 8-week post-surgery using LC-MS/MS. We found that 3 weeks after TAC, ejection fraction remained unchanged (sham: 72±5% vs TAC: 70±4%); however, myocardial carnosine and anserine levels were decreased ∼2-fold compared with sham operated hearts (**Fig. 1B**). A similar reduction (∼2-fold) was observed for 8 weeks post-TAC (**Fig. 1C**), indicating that depletion of the dipeptides plateaus sometime between 3-8 weeks.

Carnosine homeostasis in muscle is regulated via synthesis by carnosine synthase (CARNS), degradation by carnosinase (CNDP2), and cellular transport by amino acid transporters (PEPT2, TAUT, **Fig. 1A**) ^45^. To assess whether altered synthesis or transport of carnosine contributes to dipeptide depletion, we examined the expression of these enzymes and transporters at both the gene and protein levels. No changes were observed in the expression of CARNS, CNDP2, PEPT2 and TAUT at the mRNA (**Fig. 1D**) or protein levels 3 weeks post-TAC (**Fig. 1E and F**). In fact, CARNS expression in the TAC heart appears increased compared with sham despite lowered carnosine levels, although the change was not statistically significant. Overall, this suggests that reductions in carnosine levels are not due to altered synthesis, hydrolysis, or transport.

In failing hearts, there is a progressive increase in the generation of highly reactive carbonyls, such as acrolein, malondialdehyde (MDA), and 4-hydroxynonenal (HNE) which can conjugate with carnosine^18,24^. To evaluate whether aldehyde conjugation contributes to carnosine depletion, we measured the levels of the carnosine-acrolein adduct - carnosine-propanal and its reduced form, carnosine-propanol, in sham and TAC hearts using LC-MS/MS. After 3 weeks of TAC, levels of carnosine-propanal appear increased, but did not reach statistical significance (**Fig. 1G**), while carnosine propanol remained unchanged (**Fig. 1H**). In contrast, both carnosine-propanal (**Fig. 1I**) and carnosine-propanol (**Fig. 1J**) were significantly decreased 8 weeks post-TAC compared to sham hearts. Collectively, these results suggest that a decrease in myocardial histidyl dipeptides precedes overt heart failure and carnosine aldehyde conjugates increased in failing hearts might be extruded during late stages of heart failure.

### β-alanine supplementation confers cardiac protection in response to pressure overload

Following our observation that myocardial carnosine decreases prior to the onset of heart failure, we investigated whether increasing myocardial carnosine could influence the progression of heart failure. Oral administration of β-alanine, the rate limiting precursor for carnosine synthesis, enhances its synthesis and elevates intracellular levels of histidyl dipeptides^24,25^. Therefore, to test the effects of increased carnosine on heart failure progression, we placed WT C57BL/6J mice on either β-alanine in drinking water (20 g/L in drinking water) or drinking water alone for one week prior to TAC surgery. The mice were placed on β-alanine water for the entirety of the experimental period, and cardiac function was measured by echocardiography at 4 and 8 weeks after TAC (**Fig. 2A**). At 4 weeks after TAC, no significant differences in cardiac parameters were observed between β-alanine treated and water alone treated TAC mice (**Fig. 2C–H**); however, by 8 weeks post-TAC, β-alanine supplementation resulted in a significant reduction in left ventricular mass compared with mice given water alone (**Fig. 2C**). This decrease in left ventricular mass was accompanied by an improved ejection fraction (**Fig. 2B, D**), and a decrease in left ventricle inner diameter at both systole (**Fig. 2E**) and diastole (**Fig. 2F**). Additionally, β-alanine supplemented TAC mice exhibited a trend toward decreased end systolic (**Fig. 2G**) and end diastolic volumes (**Fig. 2H**), although these changes did not reach statistical significance. Collectively, these data indicate that β-alanine supplementation mitigates cardiac hypertrophy and dysfunction in response to pressure overload.

### Increased myocardial carnosine scavenges reactive aldehydes in failing hearts

Previous work has shown that lipid peroxidation products accumulate in the failing hearts and contribute to cardiac dysfunction^18,46^. Histidyl dipeptides, including carnosine, bind to these reactive aldehydes^28,30^. Because β-alanine is the precursor amino acid for carnosine, which can be further methylated to anserine^22^ or acetylated to acetylcarnosine^47^, we performed an extensive profiling of histidyl dipeptides and their aldehyde conjugates in TAC-operated hearts using LC-MS/MS to determine the impact of β-alanine treatment on the removal of toxic aldehydes. In hearts from TAC mice fed β-alanine, carnosine (*m/z* 227) and anserine (*m/z* 241) levels were significantly increased (∼10–15-fold) compared to hearts from TAC mice given water alone while N-acetylcarnosine (*m/*z 296) levels were unchanged with β-alanine supplementation (**Fig. 3A**).

**Figure 3.**
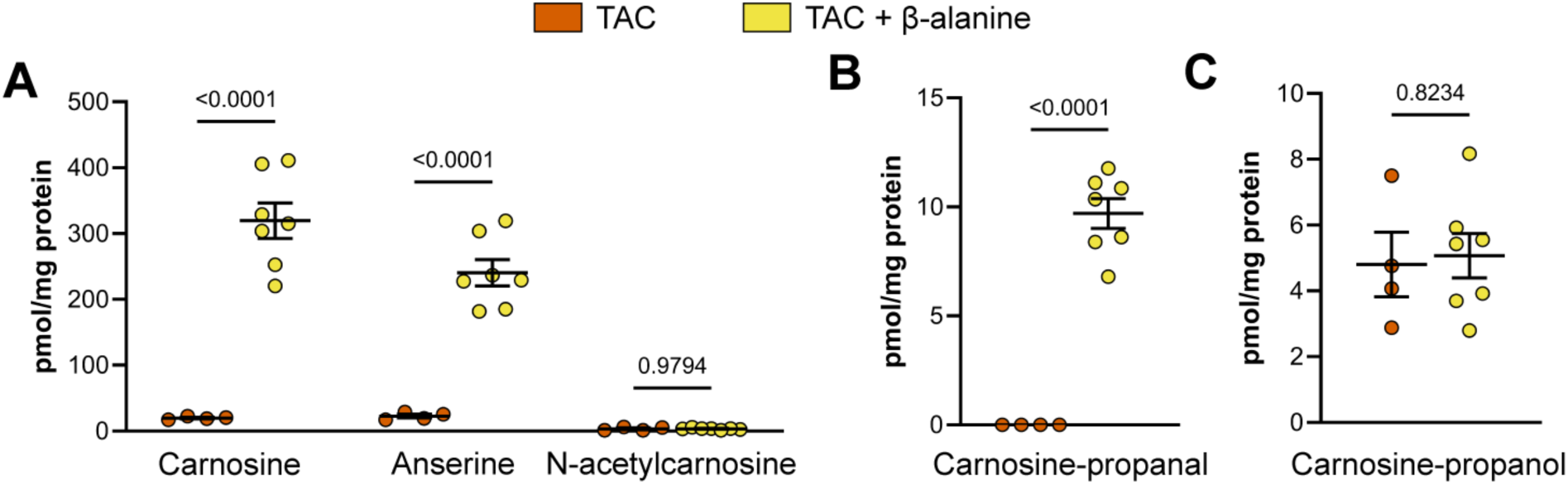
β-alanine supplementation prior to transverse aortic constriction (TAC) increases myocardial histidyl dipeptide levels and improves aldehyde quenching. Wild type (WT) C57BL/6J mice were pretreated with water alone or water supplemented with β-alanine (20 g/L) for 1 week and then subjected to TAC surgery. Treatment continued for 20 weeks. **(A)** Carnosine, anserine, and N-acetylcarnosine, **(B)** carnosine-propanal, and **(C)** carnosine-propanol levels were measured by LC-MS/MS using carnosine-d_4_ or anserine-d_4_ as an internal standard. Data shown as mean ± SEM, p-values were determined using Student’s t-test, n = 4 - 7 mice per group.

To assess the impact of β-alanine supplementation on aldehyde scavenging, we took hearts from both groups and measured conjugates of carnosine with acrolein and HNE: carnosine-propanal (*m/z* 283) and carnosine-propanol (*m/z* 285), carnosine-HNE (*m/z* 383), and conjugates of anserine with acrolein: anserine-propanal (*m/z* 297) and anserine-propanol (*m/z* 299, **Supplemental Fig. 1**). Our results show levels of carnosine-propanal conjugates were markedly increased (∼10–12-fold) in the β-alanine treated TAC hearts compared with water alone treated TAC hearts (TAC: 0 vs TAC + β-alanine: 9.7 ± 0.7 nmol/mg protein, **Fig. 3B**), while carnosine-propanol levels remained unchanged (**Fig. 3C**). No detectable levels of carnosine-HNE, anserine-propanal, or anserine-propanol were present in the TAC hearts of either group. Overall, these results indicate that increasing myocardial carnosine levels enhances the clearance of toxic aldehydes in failing hearts.

### β-alanine supplementation decreases carbonyl stress in failing heart

Having established that β-alanine supplementation increases myocardial carnosine and enhances the removal of reactive aldehydes from failing heart, we next examined whether β-alanine supplementation reduces carbonyl stress in failing heart. WT C57BL/6J mice were placed on β-alanine supplemented drinking water or drinking water alone for 1 week, followed by sham or TAC surgery with supplementation continued for an additional 4 weeks. Heart weight to tibia length ratios were significantly increased in TAC mice compared with sham controls, as expected, but this increase in heart weight was attenuated in the β-alanine treated TAC mice (**Fig. 4A**). Correspondingly, protein carbonyl content—a marker of carbonyl stress—was significantly increased in TAC hearts compared to sham and reduced in the β-alanine treated TAC hearts (**Fig. 4B**). These data suggest that carnosine-mediated aldehyde scavenging reduces carbonyl stress during heart failure.

**Figure 4.**
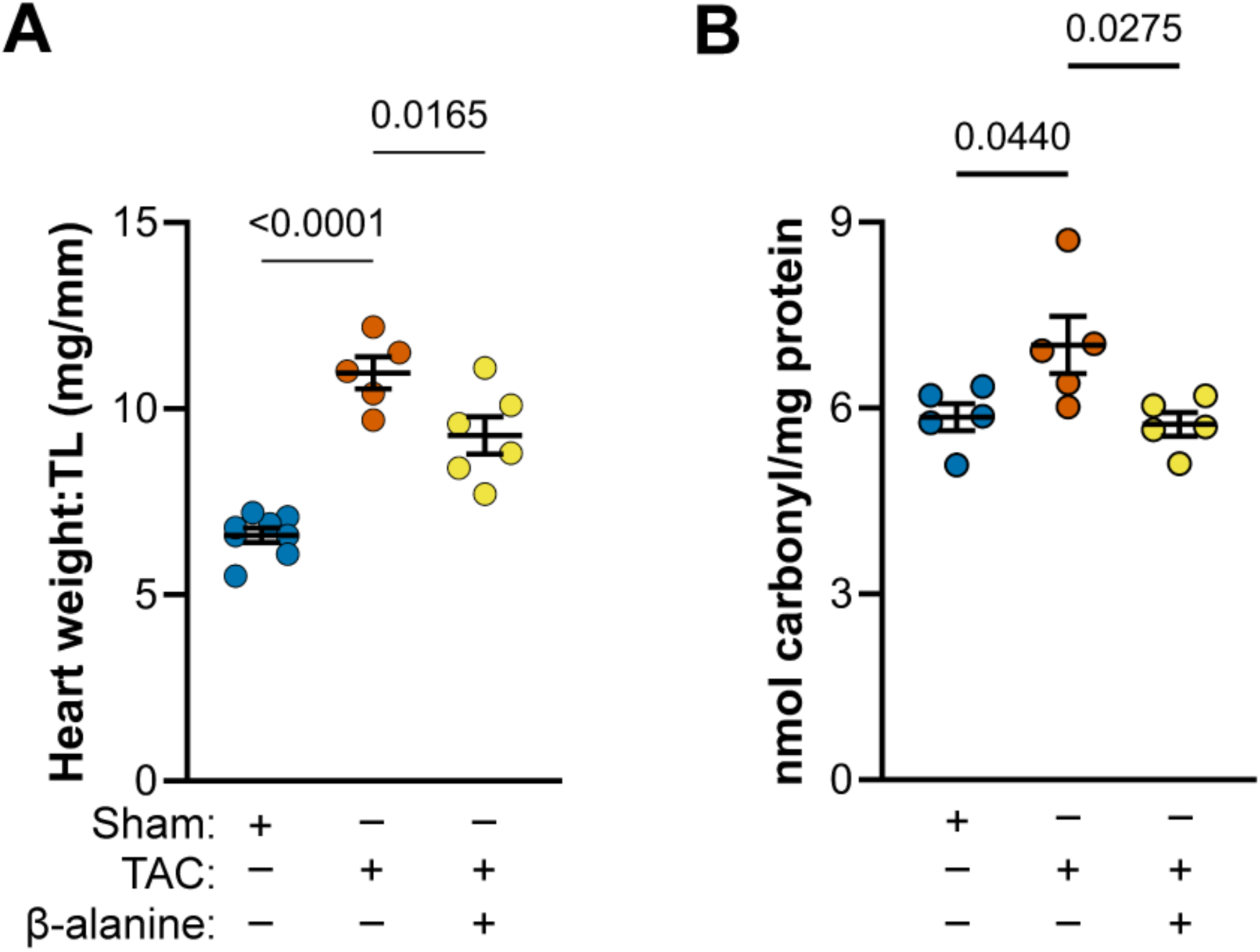
β-alanine supplementation prior to transverse aortic constriction (TAC) decreases carbonyl stress during heart failure. Male wild type C57BL/6J mice were pretreated with drinking water alone or drinking water supplemented with β-alanine (20 g/L) one week prior to TAC surgery, which was continued for 4 weeks. **(A)** Ratio of heart weight:tibia length (TL) and **(B)** carbonyl content of cardiac proteins determined by using a spectrophotometric dinitrophenylhydrazine-based assay. Data shown as mean ± SEM, p-values were determined using one-way ANOVA, n = 5 – 7 mice per group.

### Carnosine binds with aldehyde modified protein adducts

The α,β-unsaturated aldehydes possess two electrophilic reaction centers; one of the electrophilic moieties initially forms a covalent bond with the protein via either Michael adduct or Schiff base formation. Following Michael addition, the aldehyde modified protein retains the aldehyde moiety. Similarly, the Schiff base formation of the aldehyde with protein leaves a free α,β-unsaturated imine (**Fig. 5A**). The free electrophilic moiety in these aldehyde-modified proteins acts as aldehyde-modified damage associated molecular patterns (ADAMPs) that activate dendritic cells and consequently CD4^+^ T cells^14,48^. Because carnosine binds with reactive aldehydes, we next investigated whether it could bind with the free electrophilic moieties in ADAMPs forming “carnosinylated ADAMPs”. To test this, hen egg lysozyme (HEL) was incubated with HNE (1 mM) or acrolein (1 mM) to generate ADAMPs. These modified proteins were then incubated with carnosine (10 mM), sodium borohydride (NaBH_4_, 10 mM), or NaBH_4_ followed by carnosine (**Fig. 5B**). Immunoblotting with anti-HNE, anti-acrolein, and anti-carnosine antibodies showed strong immunoreactivity of ADAMPs alone with anti-acrolein and anti-HNE antibodies, while ADAMPs incubated with NaBH_4_ showed no immunoreactivity. ADAMPs incubated with carnosine had strong immunoreactivity with anti-carnosine antibodies; however, incubation of ADAMPs with sodium borohydride followed by carnosine abolished carnosine immunoreactivity (**Fig. 5C**), indicating that carnosine binds to the intact aldehyde moiety of ADAMPs.

**Figure 5.**
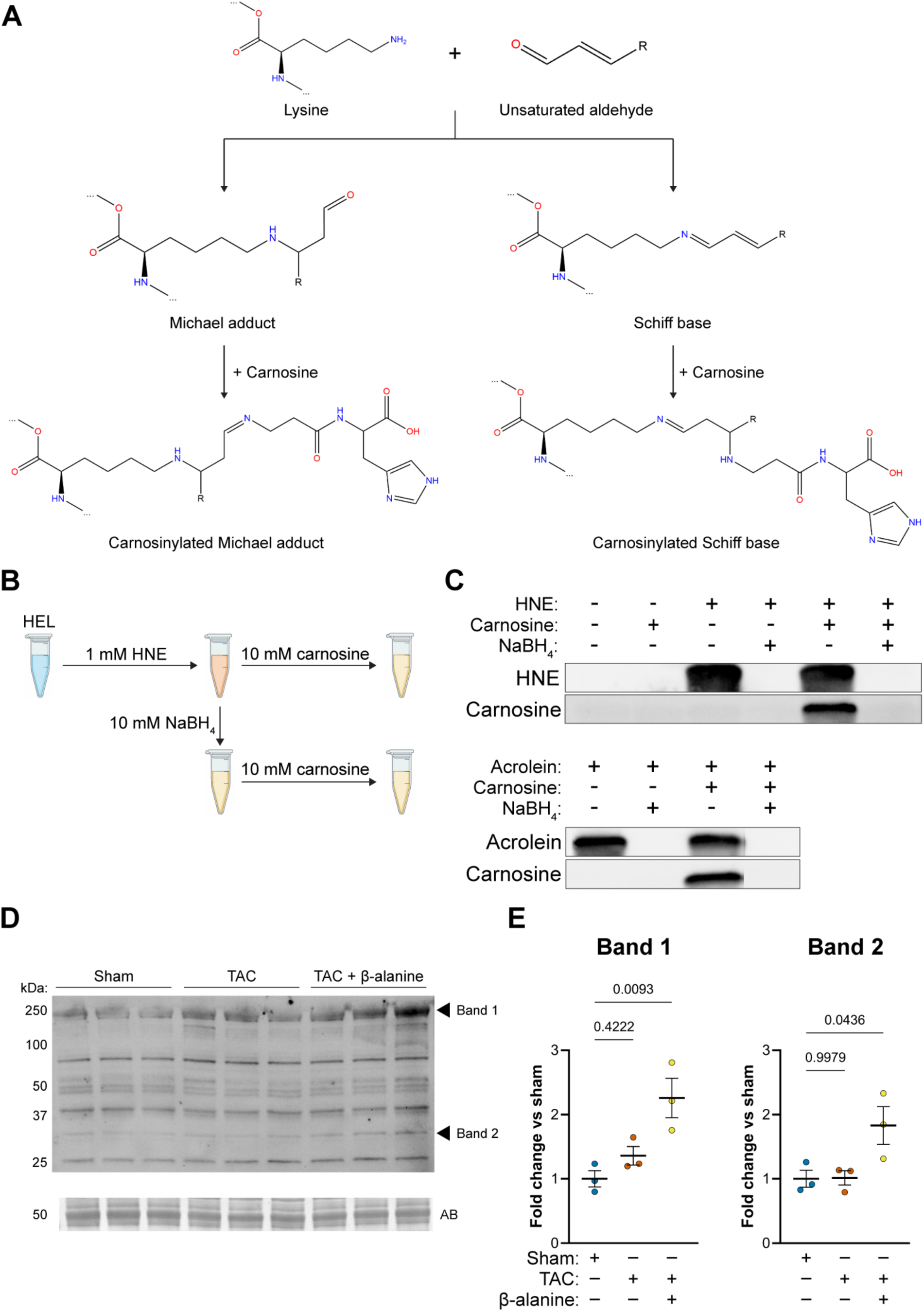
Carnosine binds to the aldehyde moiety of aldehyde-protein adducts. **(A)** Schematic demonstrating binding of reactive aldehydes with proteins either through Micheal adduct or Schiff base formation. **(B)** Hen egg lysozyme (HEL) was incubated with 1 mM 4-hydroxynonenal (HNE) or 1mM acrolein for 1 h at 37°C. The aldehyde-modified proteins were further incubated with 10 mM carnosine, reduced with 10 mM NaBH_4_, or reduced with NaBH_4_ followed by carnosine incubation (both 10 mM). **(C)** Modified HEL was immunoblotted and developed with anti-carnosine, anti-HNE and anti-acrolein antibodies. **(D)** Heart lysates from mice subjected to sham, TAC, and TAC + β-alanine supplementation were immunoblotted and developed using anti-carnosine antibody. **(E)** Intensity of bands were quantified and normalized to total protein signal using amidoblack (AB) stain. Data shown as mean ± SEM, p-values were determined using one-way ANOVA, n=3 mice in each group.

Given that ADAMPs are elevated in failing hearts, as indicated by increased carbonyl-protein adducts, and that carnosine can bind with the free aldehyde moiety of ADAMPs, we next assessed whether increasing myocardial carnosine during heart failure increases the formation of carnosinylated proteins. We found that carnosinylation of proteins was increased in the hearts of TAC mice supplemented with β-alanine when compared with both sham hearts and TAC hearts (**Fig. 5D, E**). Taken together, these results indicate that increasing myocardial carnosine during heart failure reduces cardiac carbonyl stress and promotes the formation of carnosinylated proteins.

### Carnosinylation of aldehyde modified proteins diminishes their immunogenicity

Given that carnosine binds to ADAMPs to form carnosinylated ADAMPs, we next sought to investigate whether this modification affects the immunogenicity of ADAMPs. Bone-marrow derived dendritic cells (BMDCs) were generated from WT C57BL/6J mice using GM-CSF and were subsequently incubated with 100 μg/mL unmodified hen lysozyme (HEL), MDA-modified HEL (HEL-MDA), carnosinylated HEL-MDA, or NaBH_4_ reduced HEL-MDA for 6 h. BMDC activation was measured by flow cytometry. HEL-MDA significantly upregulated MHCII and CD80 expression in BMDCs, whereas both carnosinylation and NaBH_4_ reduction abrogated these effects, implying that the binding of carnosine to the aldehyde moiety could diminish the recognition of ADAMPs by BMDCs (**Fig. 6A-D**).

**Figure 6.**
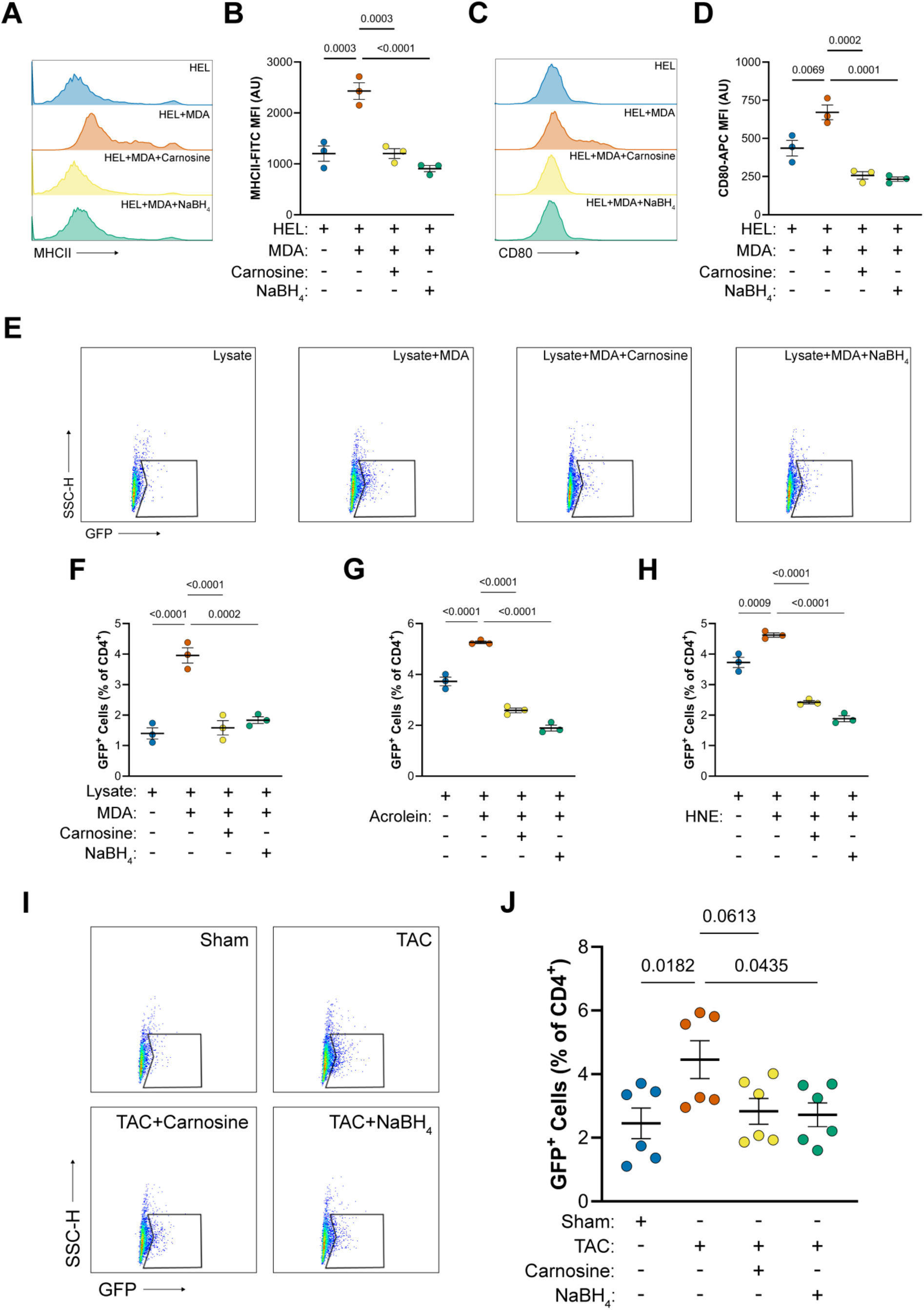
Carnosine abrogates immunogenicity of aldehyde modified cardiac proteins. **(A-D)** Bone marrow derived dendritic cells (BMDCs) were incubated with 100 μg/mL hen lysozyme (HEL), HEL-malonaldehyde, HEL-MDA incubated with carnosine, or HEL-MDA reduced by NaBH_4_ for 6 h. BMDC activation was assessed by MHCII and CD80 expression levels in CD45^+^CD11b^+^CD11c^+^ dendritic cells by flow cytometry. **(E)** Splenocytes from Nur77-GFP mice were treated with 200 μg/mL naïve cardiac lysate, lysate incubated with an aldehyde **(F**, MDA; **G**, acrolein; **H**, HNE), aldehyde-lysate incubated with carnosine, or lysate-MDA reduced with NaBH_4_. Extent of CD4^+^ T cell activation was determined by measuring GFP fluorescence in CD45^+^CD3^+^CD4^+^ cells using flow cytometry. **(I, J)** Splenocytes from Nur77-GFP mice were treated with 200 μg/mL lysate from sham mice, lysate from TAC mice, TAC lysate incubated with carnosine, or TAC lysate reduced with NaBH_4_. Extent of CD4^+^ T cell activation was then determined by measuring GFP fluorescence in CD45^+^CD3^+^CD4^+^ cells using flow cytometry. Data shown as mean ± SEM, p-values were determined using one-way ANOVA.

**Figure 7.**
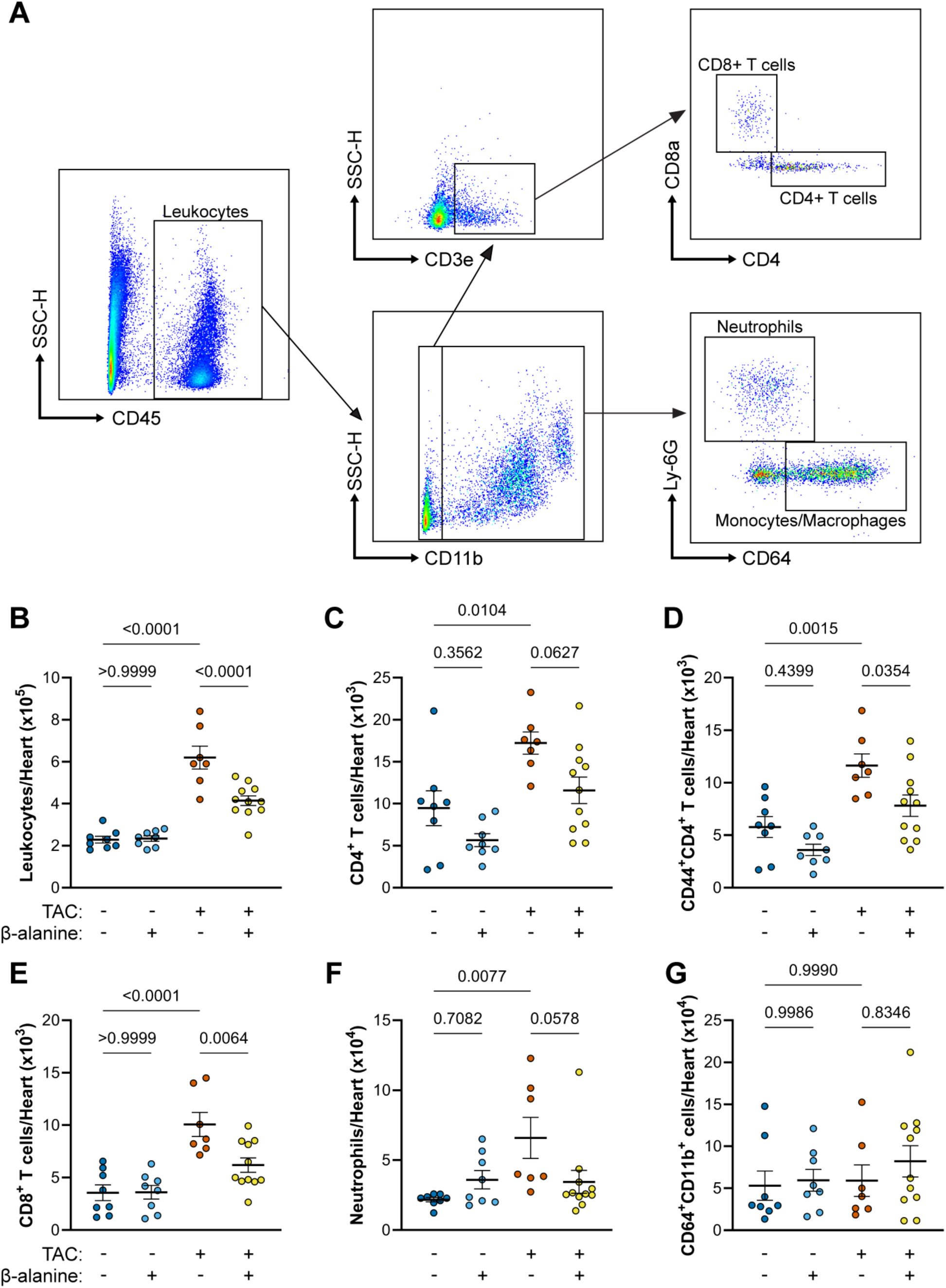
β-alanine supplementation reduces cardiac immune cell infiltration and CD4^+^ T cell activation in mice subjected to transverse aortic constriction (TAC). Male wild type C57BL/6J mice were subjected to sham or TAC surgery and given either plain drinking water or drinking water supplemented with 20 g/L β-alanine 1 week prior to surgery and continued until the end of the experiment. **(A)** 4 weeks after surgery, infiltration of **(B)** CD45^+^ leukocytes, **(C)** CD45^+^CD11b^-^CD3^+^CD4^+^ T cells, **(D)** CD45^+^CD11b^-^CD3^+^CD4^+^CD44^+^ T cells, **(E)** CD45^+^CD11b^-^CD3^+^CD8^+^ T cells, **(F)** CD45^+^CD11b^+^CD64^-^Ly6G^+^ neutrophils, and **(G)** CD45^+^CD11b^+^Ly6G^-^CD64^+^ monocytes and macrophages were determined using flow cytometry. Data shown as mean ± SEM, p-values were determined using one-way ANOVA, n = 7 - 11 mice per group.

A wide variety of reactive aldehydes are generated in failing hearts^18,46,49^. Therefore, we next investigated whether the free aldehyde moieties on cardiac proteins *per se* are able to activate CD4^+^ T cells. Further, we investigated if abrogating the antigenicity of different aldehyde moieties by carnosine would affect CD4^+^ T cell activation. To assess this, we isolated splenocytes from naïve Nur77-GFP mice, which express GFP under the control of the Nur77 promoter-a sensitive and specific readout of T cell receptor (TCR) signaling^50^. Heart lysate from naïve WT C57BL/6J mice was modified with either MDA, acrolein, or HNE (1 mM), followed by treatment with carnosine (10 mM) or NaBH_4_ (10 mM) as described in methods. Splenocytes isolated from Nur77-GFP mice were incubated with 200 μg/mL: of heart lysate alone, aldehyde modified heart lysate, aldehyde modified heart lysate incubated with carnosine, or aldehyde modified heart lysate reduced with NaBH_4_. Lysates modified with MDA, acrolein and HNE significantly increased the percentage of GFP^+^ cells within the CD4^+^ T cell gate, whereas treatment of aldehyde modified heart lysate with carnosine or NaBH_4_ eliminated the responses to these aldehyde-modified heart lysates (**Fig. 6E, H**).

Finally, to determine whether the aldehyde modified proteins generated *in vivo* during heart failure can activate CD4^+^ T cells, we incubated the splenocytes isolated from Nur77-GFP mice with 200 μg/mL: of sham heart lysate, TAC heart lysate, TAC heart lysate incubated with carnosine, or TAC heart lysate reduced with NaBH_4_. In line with our previous observations, TAC lysates significantly increased the percentage of GFP^+^ cells within the CD4^+^ T cell gate and this was abrogated in splenocytes incubated with either carnosine or reduced TAC lysates (**Fig. 6I, J**). Taken together, these data show that cardiac aldehyde modified proteins function as neoantigens capable of activating DCs and CD4^+^ T cells and that carnosine effectively masks these moieties, diminishing their neoantigenic potential.

### β-alanine supplementation inhibits CD4^+^ T cell activation and neutrophil infiltration in failing hearts

Because increasing carnosine in failing hearts scavenges reactive aldehydes, diminishes carbonyl stress, and neutralizes the neoantigenic potential of ADAMPs, we next tested whether β-alanine supplementation affects CD4^+^ T cell accumulation and activation of CD4^+^ T cells in hearts and the heart-draining mediastinal lymph nodes (mLNS) during heart failure. Male WT C57BL/6J mice were pretreated with β-alanine in drinking water or drinking water alone for 1 week, followed by sham or TAC surgery. β-alanine supplementation continued for the next 4 weeks, after which single cell suspensions from the hearts and mLNs were analyzed by FACS (**Fig. 7A**). In the hearts of TAC operated mice, the total number of leukocytes was significantly increased compared with sham hearts and this increase was significantly attenuated by β-alanine treatment (**Fig. 7B**). CD4^+^ T cells were increased in TAC hearts compared with sham and this was attenuated with β-alanine supplementation but fell short of statistical significance (**Fig. 6C**). Given that ADAMPs lead to activation of CD4^+^ T cells, next we examined the expression of CD44 on CD4^+^ T cells to discriminate between naïve and antigen-encountered cells. We found that CD4^+^CD44^+^ T cells were increased in TAC hearts and significantly reduced in β-alanine treated TAC hearts (**Fig. 6D**). Additionally, CD8^+^ T cells and neutrophils were also increased in TAC hearts and reduced upon β-alanine treatment (**Fig. 6E, F**). In contrast, monocytes and macrophages remained unchanged between the sham and TAC hearts (**Fig. 6G**). In all cell populations, β-alanine supplementation had no effect on cell numbers following sham surgery.

Finally, in mLNs, while increases were observed in the accumulation of leukocytes in TAC mice compared to sham, no significant changes were observed in dendritic cells, CD4^+^ T cells, or CD8^+^ T cells and β-alanine treatment had no effect on any of these populations (**Supplemental Fig. 2**). Taken together, these data indicate that aldehyde modified cardiac neoantigens are key drivers of CD4^+^ T cell infiltration and effector function in heart failure.

## DISCUSSION

Our data show that histidyl dipeptides are depleted before the onset of overt heart failure, likely consumed by conjugation with reactive aldehydes and supplementation with carnosine precursor amino acid β-alanine increased myocardial carnosine levels and prevented cardiac dysfunction in a pressure overload-induced model of heart failure. We demonstrate that carnosine binds not only with free aldehydes decreasing aldehyde stress but also binds with the free electrophilic group of aldehyde-modified proteins, thereby diminishing their antigenic potential and inhibiting CD4^+^ T cell activation. These findings suggest that the aldehyde-modified proteins generated in failing heart act as neoantigens, triggering innate and adaptive immune responses. Therefore, targeting aldehyde modification in the heart with carnosine either directly or through β-alanine supplementation may represent a novel and promising strategy to mitigate ADAMP-induced inflammation and associated dysfunction in the failing heart.

### Depletion of myocardial carnosine in failing hearts

Histidyl dipeptides, namely carnosine and anserine, are abundant tissues with high glycolytic activity such as the heart and skeletal muscle^24,51^, with concentrations ranging from 0.1-1 mM in healthy vertebrate hearts^24,52^. Previous work in hypertrophic and failing hearts has reported decreased myocardial carnosine following 8 weeks of transverse aortic constriction (TAC) ^44^. Targeted metabolomic analysis of serum in heart failure patients has similarly shown that carnosine as well as its constituent amino acids, β-alanine and histidine, were decreased^53^. Using the TAC model, which recapitulates the gradual transition from compensatory hypertrophy to decompensated heart failure, we found that myocardial carnosine levels decreased before the onset of overt heart failure, and this decrease in carnosine did not change further during the progression of heart failure. Although the depletion of carnosine in heart failure has been reported, the underlying mechanisms remain unclear. Carnosine homeostasis is maintained by enzymes (CARNS, CNDP2) and transporters (PHT1 and TAUT) ^45^. We found no changes in the expression of these regulators either at the onset of hypertrophy or in the failing heart, suggesting that altered synthesis or transport is likely not responsible for the observed depletion. Instead, we propose that carnosine levels are decreased as the dipeptide is consumed by binding with unsaturated aldehydes. We observed an increase in the carnosine-acrolein conjugate carnosine-propanal during early remodeling, followed by a decline in overt heart failure.

Previous work has shown that conjugates of carnosine with aldehydes are mostly excreted in urine^26,30^. Therefore, changes in the levels of these conjugates suggests these dipeptides are being consumed by binding with reactive aldehydes which might be extruded out in the urine during heart failure; however, as we did not measure the urinary levels of these conjugates in this study, further work is required to support this conclusion. Other possible contributors to the decrease in carnosine could be changes in the levels of the activities of CARNS and CNDP2, (enzymes involved in carnosine synthesis or degradation), or altered synthesis of β-alanine. Anserine levels were also decreased during the progression to heart failure, although no anserine-aldehyde conjugates were detected, suggesting binding with reactive aldehydes is unlikely to contribute to their decrease. Anserine is synthesized via enzyme carnosine methyltransferase by methylation of carnosine^54^. The concurrent reduction in both carnosine and anserine suggests that carnosine depletion may limit anserine bioavailability for methylation; however, further studies are needed to clarify whether this is due to substrate limitation or changes in enzyme activity.

### Preservation of cardiac function and decrease in aldehyde load by **β**-alanine feeding

Increasing myocardial levels of carnosine has been shown to protect against myocardial ischemia-reperfusion injury^24^. More recent work has shown that rats subjected to myocardial infarction (MI) and supplemented with β-alanine plus histidine and subjected to exercise training had improved running distance and time to exhaustion compared to exercise trained only rats; however, no benefit on cardiac function was observed by β-alanine and histidine supplementation^55^. In contrast, our study reports the benefit of β-alanine pre-treatment in ameliorating the decline in cardiac function during the late stages of heart failure. β-alanine pretreatment prevented contractile dysfunction, reduced left ventricular wall thickness and normalized heart weight/tibia length. The discrepancy with prior studies could be attributed to differences in dosage and disease models. Notably, in our study, we used 8 g/kg of β-alanine, which increased the myocardial carnosine ∼3-fold compared to the 0.25 g/kg of β-alanine used by Stefani et al ^55^, which failed to elevate myocardial carnosine levels.

Extensive evidence from humans as well as animal models of heart failure shows that there is a progressive increase in tissue levels of lipid peroxidation products, such as 4-hydroxynonenal and acrolein in failing hearts^18,56–58^. These reactive aldehydes can cause contractile and mitochondrial dysfunction in chronic heart failure^16^, and are directly correlated with impairment of left ventricular contractility in heart failure patients^59^. Consistent with these observations, we found a profound increase in the cardiac aldehyde load, characterized by increased DNPH derivatization of proteins in failing hearts. Importantly, increasing myocardial carnosine enhanced the removal of aldehydes and attenuated the aldehydic load in the context of heart failure. Collectively, these data indicate that enhancing myocardial carnosine by supplementing β-alanine removes toxic aldehydes and impedes the progression of heart failure^16^.

### Capping of aldehyde modified proteins and impact on immunogenicity

Lipid peroxidation products possess two highly electrophilic centers. Only one of the electrophilic groups can bind cysteine, histidine, and lysine residues of the proteins via formation of a Schiff base or Micheal addition followed by Amadori rearrangement, generating a multitude of aldehyde modified protein adducts ^12,60^. These aldehydes modified proteins are not end products; they are intermediates that possess the potential to bind with free thiols via a Michael type addition forming glutathiolated protein adducts^12^. Because the aldehyde-modified proteins still retain at least one electrophilic moiety, and carnosine binds to electrophiles through the terminal or imidazole amines, we expected carnosine to bind and carnosinylate the free aldehyde moiety of aldehyde modified proteins. Indeed, our observations show that the incubation of HNE and acrolein modified HEL with carnosine increased the immunoreactivity with anti-carnosine antibody, whereas this immunoreactivity was abrogated in the aldehyde modified HEL reduced with sodium borohydride. These findings suggest that carnosine is incorporated into aldehyde modified proteins by binding with a free electrophilic moiety. Previously, we reported that carnosinylated proteins accumulate in aged and ischemic heart tissues^30^; however, to the best of our knowledge, whether the carnosinylation of aldehyde modified proteins could be influenced by heart failure conditions or altered by increasing carnosine levels has not been reported. The present study reveals a previously unrecognized mechanism by which increasing myocardial carnosine in the failing heart modulates protein carnosinylation and could potentially diminish the neoantigenic potential of aldehyde modified proteins in failing hearts.

### Aldehyde-modified damage associated molecular patterns (ADAMPs) and the role of carnosine in modulating adaptive immunity

Extensive evidence shows that dying cardiomyocytes from failing hearts release multiple damage associated molecular patterns (DAMPs), which are recognized by pattern recognition receptors (PRR) and activate innate and adaptive immune responses; however, many of these DAMPs have both protective and harmful effects. For example, intracellular S100 proteins, heat shock protein 60 (HSP60), high-mobility group box 1 (HMGB1), mitochondrial DNA, and isolevuglandin modified proteins released from failing hearts are all recognized as DAMPs and contribute to the development of heart failure^9,46,61,62^. On the other hand, HSP60 is essential for normal mitochondrial and cardiomyocyte homeostasis^55^. Similarly, differential effects of HMGB1 inhibition were observed during ischemia-reperfusion injury in heart^8,63,64^. Administration of HMGB1 has also been shown to improve ejection fraction post myocardial infarction^65^. This duality highlights the need to differentiate harmful DAMPs from DAMPs which may have benign or protective functions. Aldehyde modified protein adducts have long been identified as triggers of innate and adaptive immune responses in several oxidative stress associated diseases, including atherosclerosis and exposure to environmental contaminants^66,67^. The free electrophilic group of aldehyde-modified proteins causes them to act as neoantigens, or aldehyde modified damage associated molecular patterns (ADAMPs). The ADAMPs are taken up and presented by dendritic cells, leading to the activation of T-cells^66^.

The results of our present study show that model ADAMPs (HEL-MDA) activate DCs, as determined by increased expression of MHCII and co-stimulatory molecules CD80 and CD86. We further found that the incubation of HEL-MDA with whole splenocyte populations isolated from Nur77^GFP^ transgenic T cell reporter mice (Tg) mice increase GFP expression in CD4^+^ T cells, whereas reduction of the aldehyde moiety by sodium borohydride dampens DC and CD4^+^ T cell activation. These findings are aligned with previous observations that the aldehyde moiety specifically causes aldehyde-modified proteins to behave as immunogenic antigens^14^. Our results show that aldehyde-modified heart lysate, which mimics failing hearts loaded with ADAMPs, and homogenate from hypertrophic TAC hearts upregulated GFP expression in CD4^+^ T cells in splenocyte populations from Nur 77^GFP^ mice. These findings support our model that ADAMPs generated in failing hearts are presented by DCs, which ultimately activate CD4^+^ T cells during heart failure. Our findings provide an assessment of pathophysiological significance of protein carnosinylation. Capping of the free aldehyde moiety by carnosine of HEL-MDA, aldehyde modified heart lysate, or ADAMP-containing heart failure lysate diminished the GFP expression of CD4^+^ T cells from Nur77^GFP^ mice. However, we did not identify the specific ADAMPs generated in failing hearts that trigger immune responses, and further studies are needed to identify the specific epitopes on these ADAMPs which trigger inflammation. Nevertheless, our data strongly suggest that ADAMPs generated in failing hearts act as neoantigens that activate DCs and ultimately CD4^+^ T cells. Importantly, we report that posttranslational modification of ADAMPs by carnosine diminishes signaling between the DCs and CD4^+^ T cells. Therefore, this posttranslational modification of ADAMPs by carnosine could be an effective approach to dampen the DCs and CD4^+^ T cells activation in failing hearts.

### Diminished CD4^+^ T cell activation and neutrophil infiltration in failing hearts by **β**-alanine feeding

Several studies using the TAC model of heart failure have demonstrated that DCs, neutrophils, and CD4^+^ T cells infiltrate the left ventricle (LV) and contribute to the pathogenesis of heart failure^68–71^. DCs also accumulate in the mediastinal lymph nodes (mLNs) that drain the heart^46^. Within the LV, these accumulated DCs promote adverse remodeling by presenting DAMPs to and activating CD4^+^ T cells, which then infiltrate the myocardium. Notably, depletion of bone marrow derived DCs significantly attenuates left ventricular fibrosis and hypertrophy in response to TAC^72^. Similarly, studies using T cell deficient mice show recombinase-activating gene null (Rag2^-/-^)^68^, T cell receptor null (Tcr^-/-^)^69^ and major histocompatibility complex II null (MHCII^-/-^)^68^ mice do not develop adverse cardiac remodeling in the onset of TAC. These findings collectively suggest a critical role for DCs and T cell interaction in mediating cardiac inflammation and tissue remodeling.

Further, oxidized proteins generated in response to TAC or hypertension serve as antigens that are presented by DCs, leading to T-cell activation^46,73^. We found that TAC promotes the accumulation of DCs and CD4^+^ T cells in the mLNs and heart. This accumulation of DCs and CD4^+^ T cells coincides with the onset of aldehyde stress, implicating ADAMPs as potential triggers for DCs and CD4^+^ T cell activation. Our results show that the population of effector T-cells (CD4^+^ CD44^+^ T cells) significantly increases in response to TAC and that treatment with β-alanine diminishes this accumulation. Furthermore, TAC induced neutrophil infiltration was decreased by β-alanine supplementation. Previous studies have shown that lipid peroxidation products promote neutrophil chemotaxis^74^ and that CD4^+^ T cells increase neutrophil survival time^75^. Our findings that β-alanine supplementation increases sequestration of lipid peroxidation products and reduces activated CD4^+^T cells in failing hearts suggests a combinatorial effect contributing to neutrophil depletion. Together these findings support the hypothesis that increasing myocardial carnosine levels through β-alanine supplementation facilitates the sequestering of reactive aldehydes and carnosinylation of ADAMPs in the failing heart. This process likely reduces aldehyde stress, diminishes neoantigenicity of ADAMPs, and consequently alleviates the maladaptive immune response.

## Conclusions

Although oxidative stress is a well-established contributor to the pathogenesis of heart failure, treatment with antioxidants has limited clinical success in treating cardiovascular disease^76,77^. Our data suggest that increasing myocardial carnosine can attenuate heart failure through binding with free aldehydes, the downstream effectors of oxidative stress. Further, carnosine binds with the aldehyde moiety of ADAMPs, offering a novel strategy to dampen immune activation during heart failure. Finally, myocardial carnosine levels can be safely and effectively increased through oral administration of β-alanine^24^, and thus carnosine represents a novel intervention to reduce inflammation and preserve cardiac function during heart failure.

### Study limitations

β-alanine supplementation increases carnosine levels in the brain and skeletal muscle, yet whether the effects of β-alanine on heart failure outcomes are cardiocentric still needs to be determined. In addition, the specific ADAMPs and the core peptides of these ADAMPs which are immunogenic still need to be identified and the extent of their immunogenicity has yet to be determined.

## PERSPECTIVES

### Competency in medical knowledge

<u>Inflammation and formation of aldehyde modified protein adducts the downstream effectors of oxidative stress</u> is a cardinal feature of heart failure. In the heart, there are present small histidyl dipeptides, such as carnosine which binds with toxic aldehydes. Our findings show levels of these dipeptides are depleted, supplementation with β-alanine, the precursor for carnosine increase its myocardial levels and preserves cardiac function during heart failure. Mechanistically, increasing myocardial carnosine levels diminishes aldehyde stress, antigenicity of aldehyde modified protein adducts, and subsequently the activation of CD4+ T cells in failing hearts.

### Translational outlook

No effective drug is currently available for heart failure patients to prevent inflammation. Small histidyl dipeptides, such as carnosine present in the heart, can bind with the toxic lipid peroxidation products, aldehyde modified protein adducts and concomitantly diminish aldehyde stress and inflammatory responses. Since carnosine and β-alanine are food supplements, safe for human consumption and there levels in different tissues especially heart can be enhanced by supplementation, thus increasing the myocardial levels of carnosine can be beneficial for heart failure patients. Our results showing that β-alanine supplementation prevents cardiac dysfunction, lays the foundation for testing the benefits of supplementing to β-alanine to heart failure patients.

## ACKNOWLEDGEMENTS

We acknowledge helpful discussions and the expert technical assistance of members of our laboratories and others in the Center for Cardiometabolic Science. In addition, we acknowledge the various cores that made this work possible, including the Physiology, Microscopy and Pathology, and Flow Cytometry Cores in the Center and the Mass Spectrometry Core in the Envirome Institute.

## Contributions

BD: Designed and performed experiments; analyzed and interpreted data; provided funding; wrote and revised manuscript

MC, IJ: Performed experiments; analyzed data

DH, KB, YN: Designed and performed experiments; interpreted data

JKS: Assisted experimental design and execution

TM, MW: Designed experiments; interpreted data; revised manuscript

AB: Provided funding; revised manuscript

SPJ: Designed experiments; interpreted data; wrote and revised manuscript

SPB: Conceived project; designed experiments; interpreted data; provided funding; wrote and revised manuscript

## ABBREVIATIONS AND ACRONYMS

ADAMPs: Aldehyde-modified damage associated molecular patterns
BMDCs: Bone marrow-derived dendritic cells
CARNS: Carnosine synthase
GFP: Green fluorescent protein
GM-CSF: Granulocyte/monocyte colony stimulating factor
LC-MS/MS: Liquid chromatography with tandem mass spectrometry LVIDd – Left ventricular internal diameter at diastole
LVIDs: Left ventricular internal diameter at systole
mLNs: Mediastinal lymph nodes
TAC: Transverse aortic constriction

---

**Supplemental Figure 1.** V**e**rification **of LC-MS chromatograms.** Heart lysates were prepared in 10 mM HCl containing 5 μM carnosine-d4 and 5 μM anserine-d4 as internal standards. Samples were injected into a Waters ACQUITY UPLV H-Class system and separated using a HILIC protocol. Retention times and structures are shown for **(A)** carnosine, **(B)** anserine, **(C)** N-acetylcarnosine, **(D)** carnosine-propanal, and **(E)** carnosine-propanol.

**Supplemental Figure 2**. Leukocytes infiltrate mLNs following TAC but are unaffected by β-alanine supplementation. Male Wild type C57BL/6J mice were subjected to sham or TAC surgery and given either plain drinking water or drinking water supplemented with 20 g/L β-alanine 1 week prior to surgery and continued until the end of the experiment. 4 weeks after surgery, single cell suspensions were made from mediastinal lymph nodes (mLNs). (A) Total CD45+ leukocytes, (B) CD45+CD11b-CD3+CD4+ T cells and (C) CD45+CD11b-CD3+CD8+ T cells. (D) Dendritic cells were counted in the mLNs by gating on CD45+CD11b+CD11c+ populations and were evaluated for markers of (E) maturation (MHCII) and (F, G) activation (CD80 and CD86).

