## Supplemental Appendix for "Carnosinylation of Cardiac Antigens Attenuates Immunogenic Responses and Improves Function in Failing Hearts"

| Supplemental Methods | p. 2-3 |
| --- | --- |
| Supplemental Figures | p. 4-5 |

**SUPPLEMENTAL METHODS**

**Table I. Fluorescent antibodies for flow cytometry**

| Target | Fluorophore | Clone | Supplier | Cat. # |
| --- | --- | --- | --- | --- |
| CD45 | PE/Cy7 | 30-F11 | Biolegend | 103114 |
| CD11b | BV 510 | M1/70 | Biolegend | 101245 |
| CD11b | PE/Dazzle 594 | M1/70 | Biolegend | 101255 |
| CD11c | PE | N418 | Biolegend | 117307 |
| CD80 | APC | 16-10A1 | Biolegend | 104713 |
| CD86 | BV 421 | GL-1 | Biolegend | 105032 |
| MHCII (I-A/I-E) | FITC | AF6-120.1 | Biolegend | 107605 |
| Ly6G | APC | 1A8 | Biolegend | 127613 |
| CD64 | PE | X54-5/7.1 | Biolegend | 139303 |
| CD3e | RealBlue 705 | 145-2C11 | BD Biosciences | 570560 |
| CD4 | BV 750 | GK1.5 | Biolegend | 100467 |
| CD4 | APC | GK1.5 | Biolegend | 100412 |
| CD8a | BV 650 | 53-6.7 | BD Biosciences | 563234 |
| CD44 | PE/Cy5 | IM7 | Biolegend | 103010 |

**Table II. RT-qPCR primers:** All primers were purchased from Integrated DNA Technologies and primer specificity was verified using NCBI Primer-BLAST. Sequences used were as follows:

| Gene | Forward (5’ → 3’) | Reverse (5’ → 3’) |
| --- | --- | --- |
| *Carns1* | TGATAGGCCCCTACTGAGTAAGGT | TCAGTGTCCTTGGCAGGGTAT |
| *Cndp2* | GGAGATACCACTTCCTCCCATCT | CGTCCAGGTGCCCGTAAAT |
| *Pept2* | TGGCTGGGAAAATTCAAGACA | ATGGCACCCAAAGACTTGAATAC |
| *Taut* | TGGCCGACAGCATTCCA | GCCTTCTCTAAGGTGCCTTCCT |
| *Hprt* | AGGACCTCTCGAAGTGTTGG | AGGGCATATCCAACAACAAAC |

**SUPPLEMENTAL FIGURES**

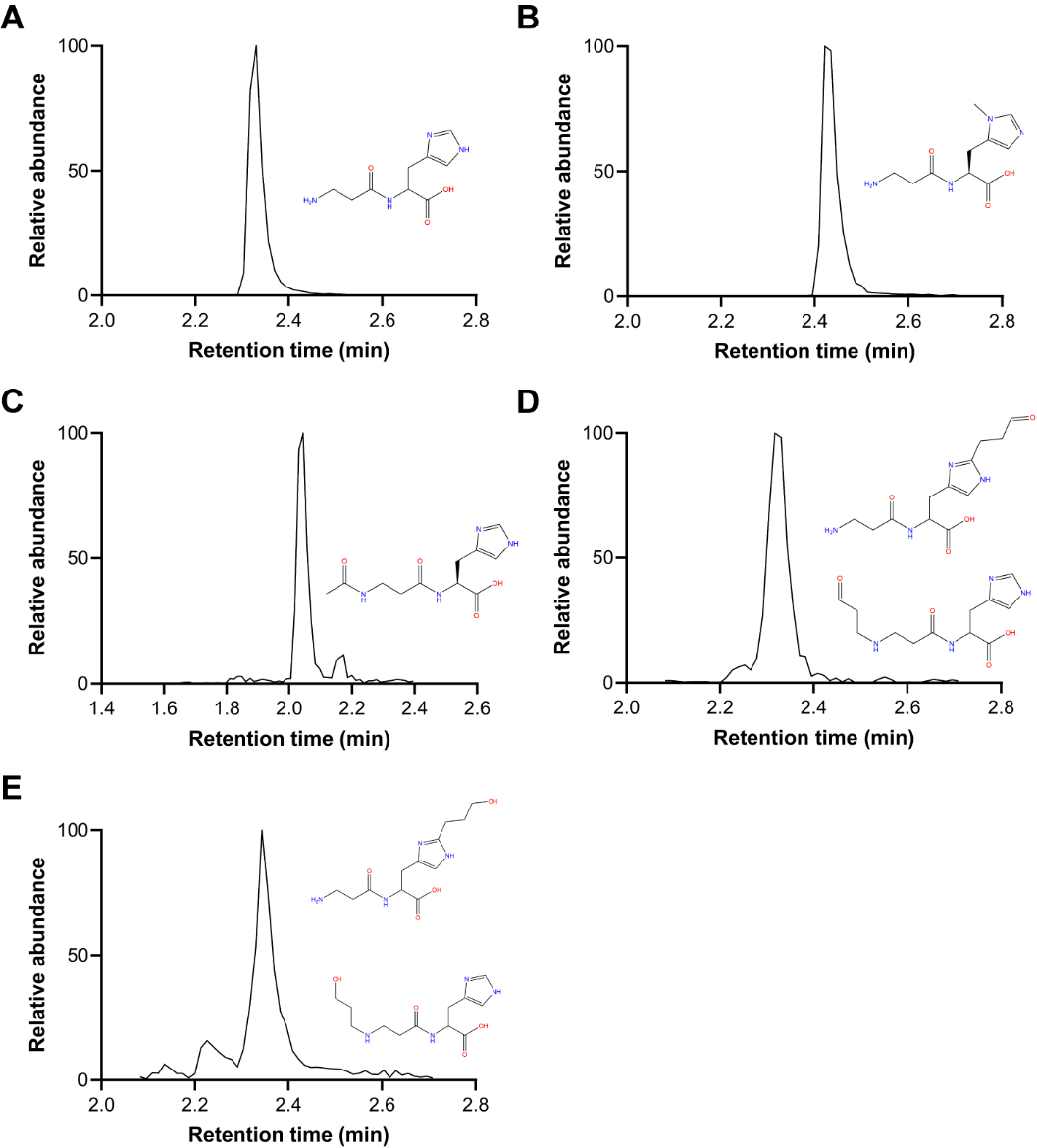

**Supplemental Figure 1. Verification of LC-MS chromatograms.** Heart lysates were prepared in 10 mM HCl containing 5 μM carnosine-d4 and 5 μM anserine-d4 as internal standards. Samples were injected into a Waters ACQUITY UPLV H-Class system and separated using a HILIC protocol. Retention times and structures are shown for **(A)** carnosine, **(B)** anserine, **(C)** N-acetylcarnosine, **(D)** carnosine-propanal, and **(E)** carnosine-propanol.

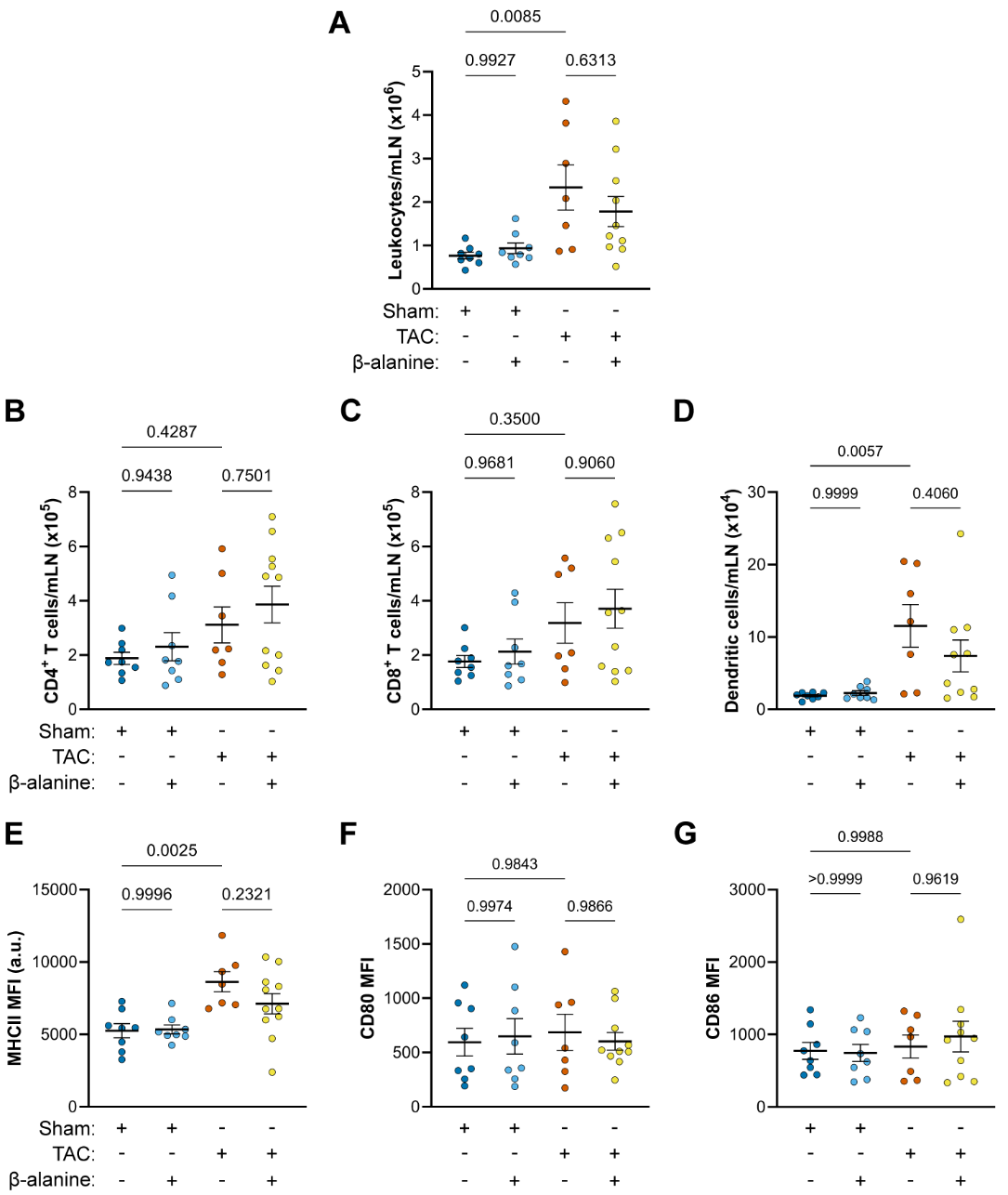

**Supplemental Figure 2. Leukocytes infiltrate mLNs following TAC but are unaffected by β-alanine supplementation.** Male Wild type C57BL/6J mice were subjected to sham or TAC surgery and given either plain drinking water or drinking water supplemented with 20 g/L β-alanine 1 week prior to surgery and continued until the end of the experiment. 4 weeks after surgery, single cell suspensions were made from mediastinal lymph nodes (mLNs). **(A)** Total CD45^+^ leukocytes, (**B**) CD45^+^CD11b^-^CD3^+^CD4^+^ T cells and (**C**) CD45^+^CD11b^-^CD3^+^CD8^+^ T cells. **(D)** Dendritic cells were counted in the mLNs by gating on CD45^+^CD11b^+^CD11c^+^ populations and were evaluated for markers of **(E)** maturation (MHCII) and **(F, G)** activation (CD80 and CD86).
